## Supplemental Figures for "Roles of P-body factors in *Candida albicans* filamentation and stress response"

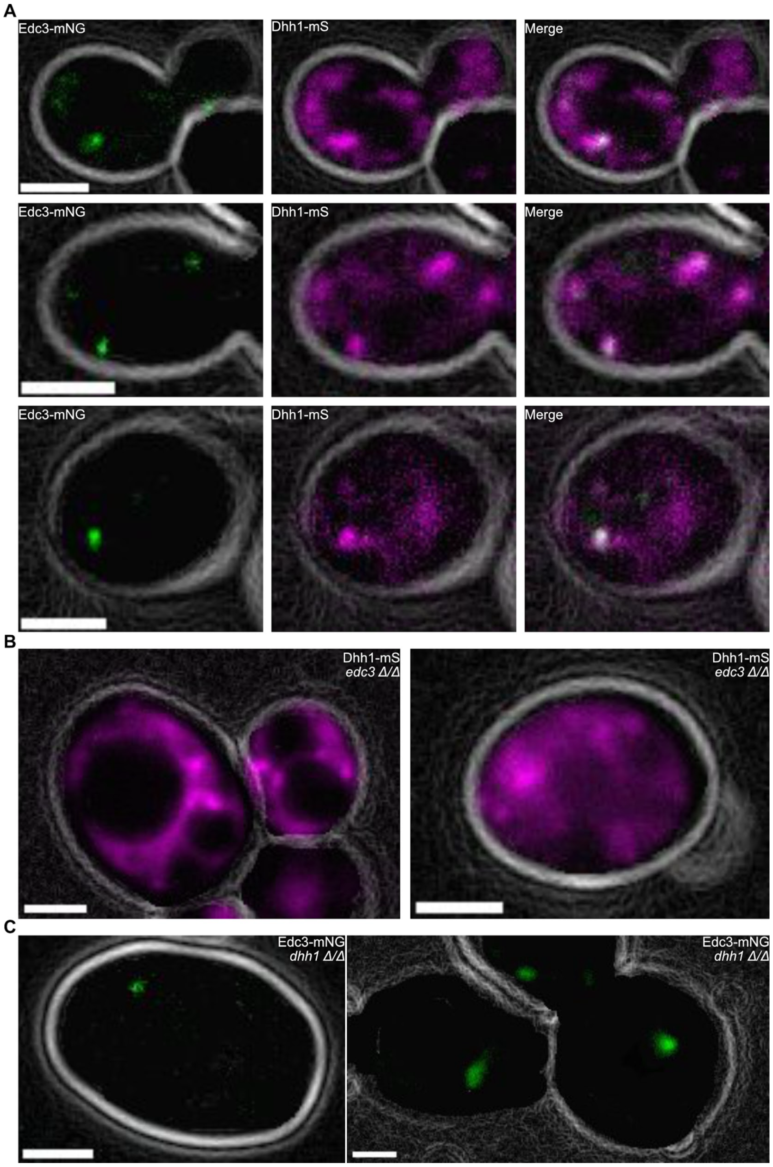


**Fig S1. Replicates of PBs condensation during heat stress in SC5214. A.** PB factors (Edc3-mNG and DHH1-mS) co-localized during acute heat shock (10 minutes at 46°C in CM) confirming their identity as PBs. Scale bars = 2μm. **B.** Dhh1-mS condensed in the absence of Edc3 in response to acute heat shock (10 minutes at 46°C in CM). Scale bars = 2μm. **C.** Edc3-mNG condensed in the absence of Dhh1 response to acute heat shock (10 minutes at 46°C in CM). Scale bars = 5μm.

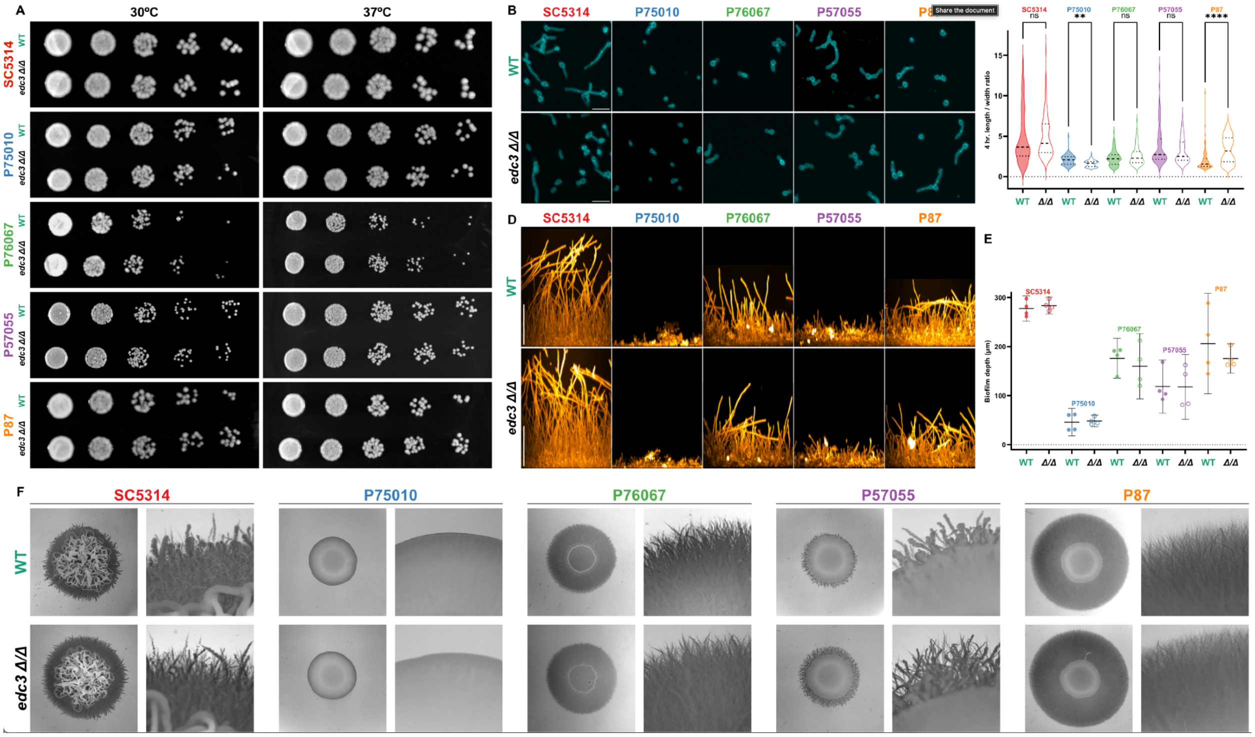


**Fig S2. Absence of *EDC3* does not impair growth or hyphal morphology in diverse strains.** WT and *edc3Δ/Δ* strains were subjected to growth, filamentation, biofilm, and spider plate assays. **A.** 24-hour spot plate assay of *C. albicans* WT and *edc3Δ/Δ* shows *EDC3* is not required for WT growth in all 5 strains. **B.** *edc3Δ/Δ* strains were subjected to a planktonic 4-hour filamentation assay (37°C, RPMI + 10% serum) and did not display gross inhibition of filamentation compared to WT. Scale bars = 20µm. **C.** After 4-hours of filamentation, *edc3Δ/Δ* did not have grossly impaired germ tube development. Only *edc3Δ/Δ* P75010 was significantly less hyphal than WT (N =100 cells, ****P < 0.0001, **P < 0.01, Kruskal-Wallis tests). **D.** Side-view of WT and *edc3Δ/Δ* strains after a 24-hour biofilm assay (37°C, RPMI + 10% serum). Scale bar = 100µm. **E.** Depth of 4 replicate biofilms. 95% CI. **F.** Colonies grown on spider plates for 6 days at 37°C. 6X and 25X magnification.

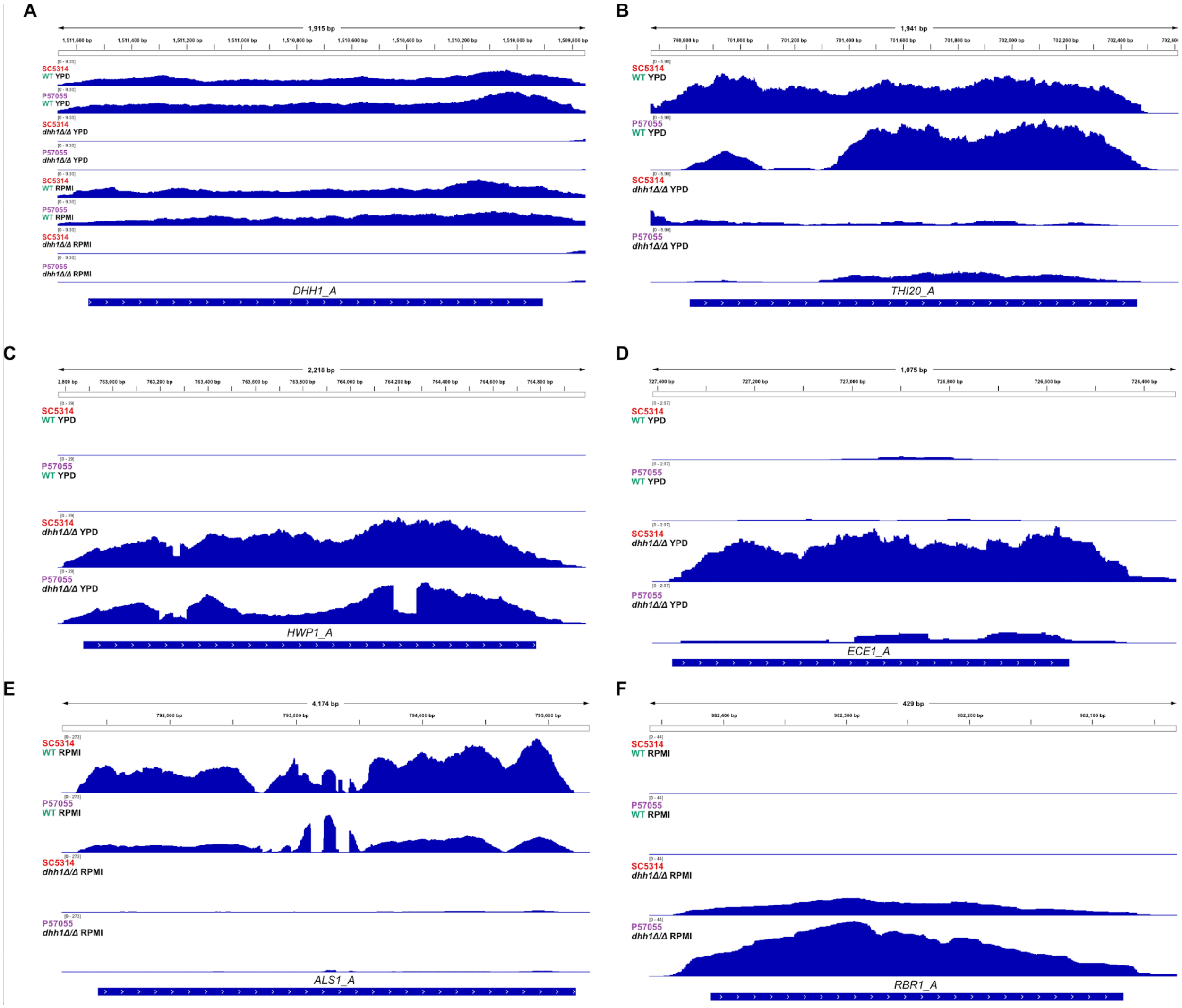


**Fig S3. Normalized IGV tracks for SC5314 and P57055 WT and *dhh1Δ/Δ* strains.** Tracks normalized by cpm were examined to verify deletions and differentially expressed genes **A.** *DHH1* transcripts are present in WT backgrounds and absent from *dhh1Δ/Δ*. **B.** In YPD putative trifunctional thiamine biosynthesis enzyme *THI20* is downregulated in both *dhh1Δ/Δ* strains. **C.** Hyphal wall protein (*HWP1*) transcripts are upregulated in *dhh1Δ/Δ* compared to WT under yeast-form growth conditions. **D.** Interestingly, in YPD hyphal associated candidalysin (*ECE1*) is upregulated in SC5341 *dhh1Δ/Δ*, and to a lesser extent in P57055 *dhh1Δ/Δ*. **E.** Hyphal adhesin (*ALS1*) is downregulated in both *dhh1Δ/Δ* strains in RPMI, **F.** while GPI-anchored cell wall protein (*RBR1*) is upregulated in both *dhh1Δ/Δ* strains under these conditions.

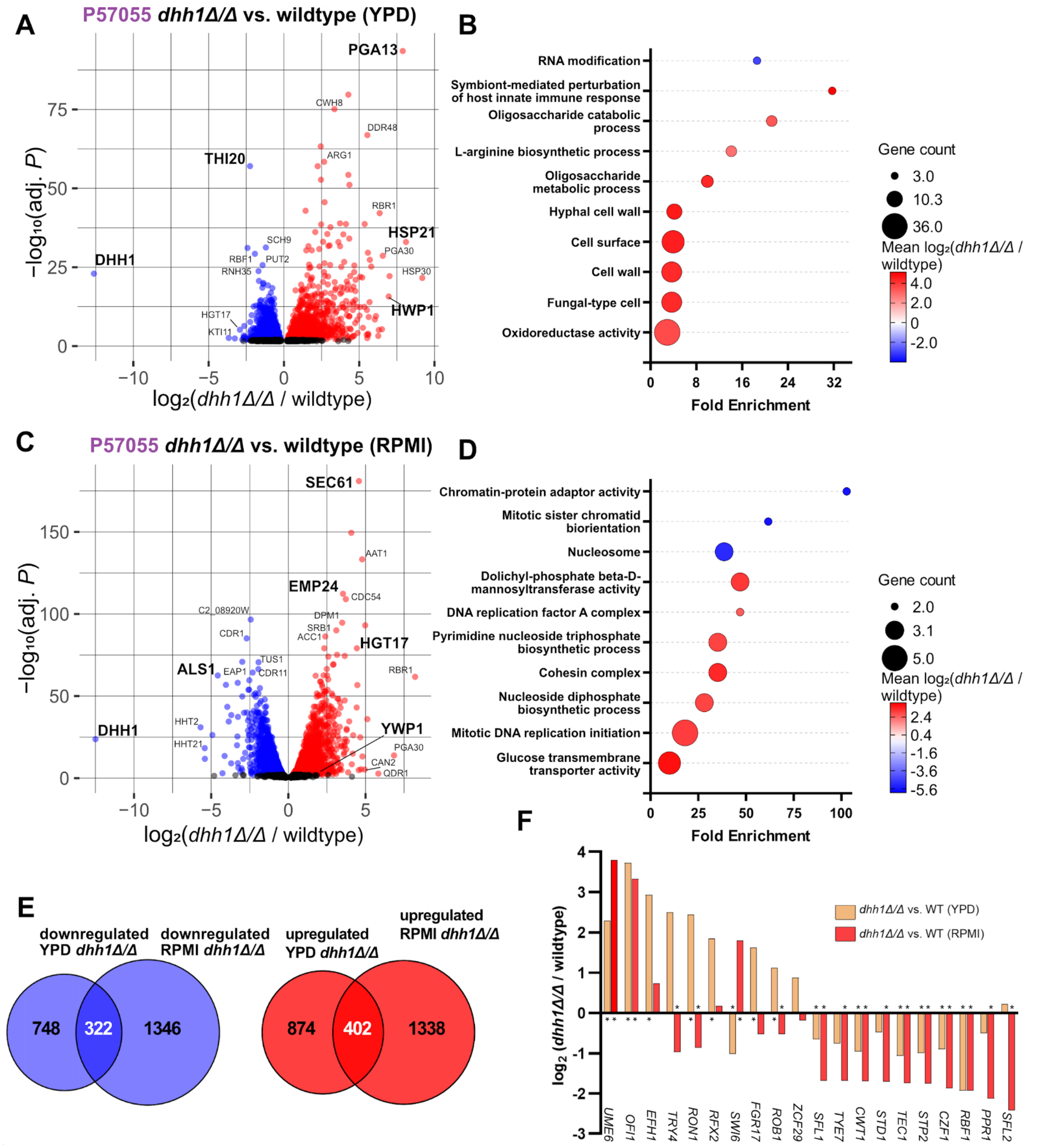


**Fig S4. RNA-seq exposed differential gene expression in P57055 *dhh1Δ/Δ* grown under YPD and RPMI conditions. A.** Volcano plot depicts genes significantly up (red) and down (blue) -regulated during yeast-form growth compared to wildtype (Padj < 0.01). Genes associated with filamentation stress were prominently upregulated (Table S5). **B.** GO terms enriched in *dhh1Δ/Δ* YPD include transport and cell wall terms (Padj < 0.1). in. **C.** Volcano plot depicts gene expression differences in *dhh1Δ/Δ* vs wildtype during early hyphal growth (Table S6, Padj < 0.01). **D.** GO terms enriched in *dhh1Δ/Δ* RPMI are related to chromatin, replication and biosynthesis (Padj < 0.1). **E.** A significant number of transcripts were downregulated (P = 0.041), and upregulated (P = 0.0178), respectively, in *dhh1Δ/Δ* in YPD and RPMI conditions (Hypergeometric tests). However, most differentially expressed genes were environmentally specific. **F.** Differential expression of transcription factors regulating morphogenesis in *dhh1Δ/Δ* in YPD and RPMI. (* indicates Padj < 0.01; DEseq2).

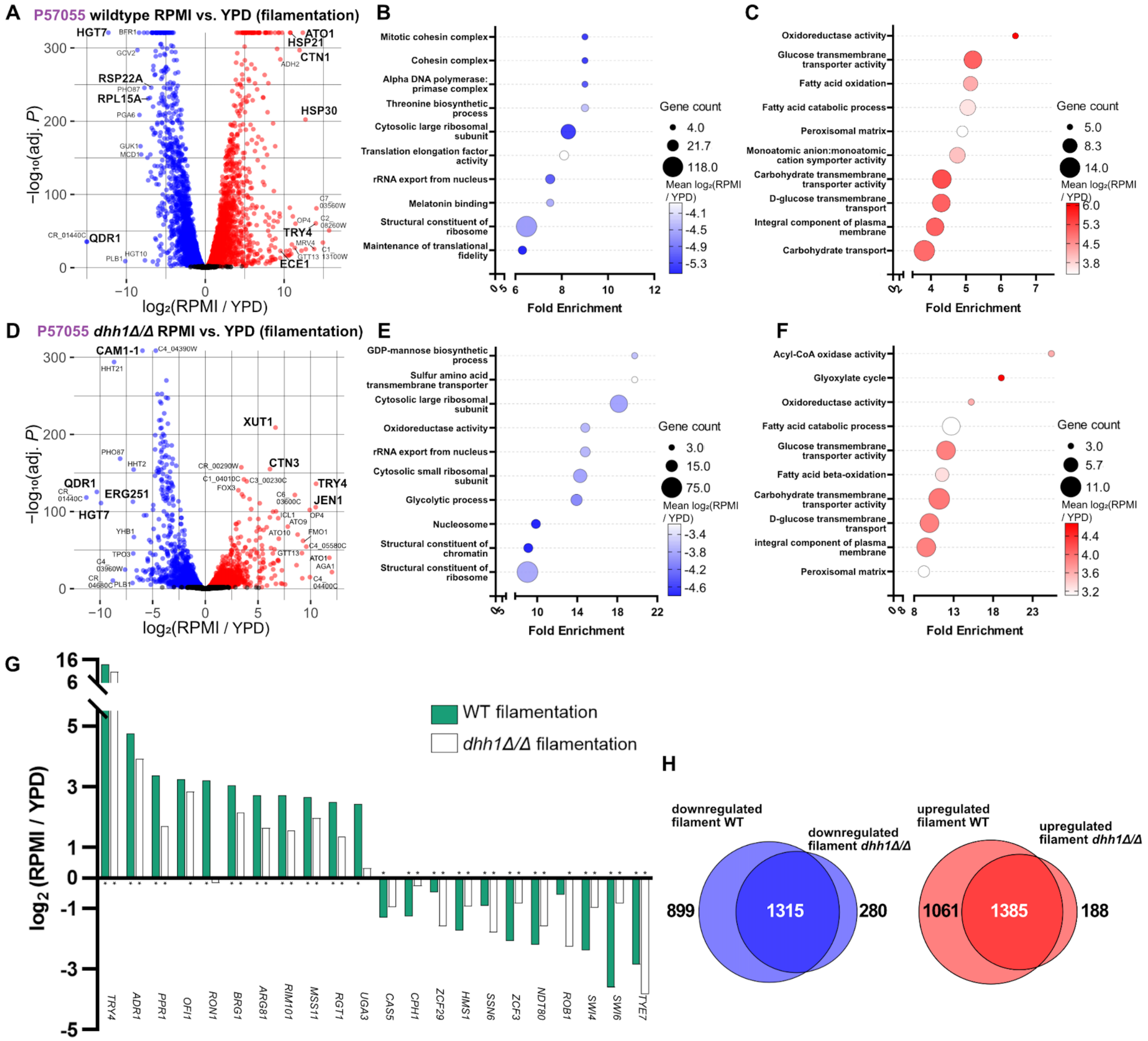


**Fig S5. RNA-seq reveals transcriptome differences between WT and *dhh1Δ/Δ* during filamentation.** **A.** Volcano plots comparing the transcriptomes of WT P57055 grown in YPD and RPMI demonstrate extensive remodeling during filamentation (Table S7, Padj < 0.01). **B.** Expected GO terms were downregulated, and **C.** upregulated during WT filamentation (Padj < 0.1). **D.** Comparing the transcriptome of *dhh1Δ/Δ* grown in YPD and RPMI showed transcriptome remodeling during filamentation (Table S8, Padj < 0.01). **E.** Similar GO terms were downregulated, and **F.** upregulated in filamenting *dhh1Δ/Δ* and WT (Padj < 0.1). Unlike WT, *dhh1Δ/Δ* upregulated terms related to lipid catabolism during filamentation. **G.** Transcription factors regulating morphogenesis were differentially expressed during filamentation in WT and *dhh1Δ/Δ* (* indicates Padj < 0.01; DEseq2). **H.** Genes differentially expressed during filamentation overlapped between WT and *dhh1Δ/Δ* strains. Most transcripts differentially expressed during *dhh1Δ/Δ* filamentation are subsets of the WT remodeling response. A significant number of transcripts were down- (P = 0) or upregulated (P = 0) in both filamenting *dhh1Δ/Δ* and wild type (Hypergeometric tests).

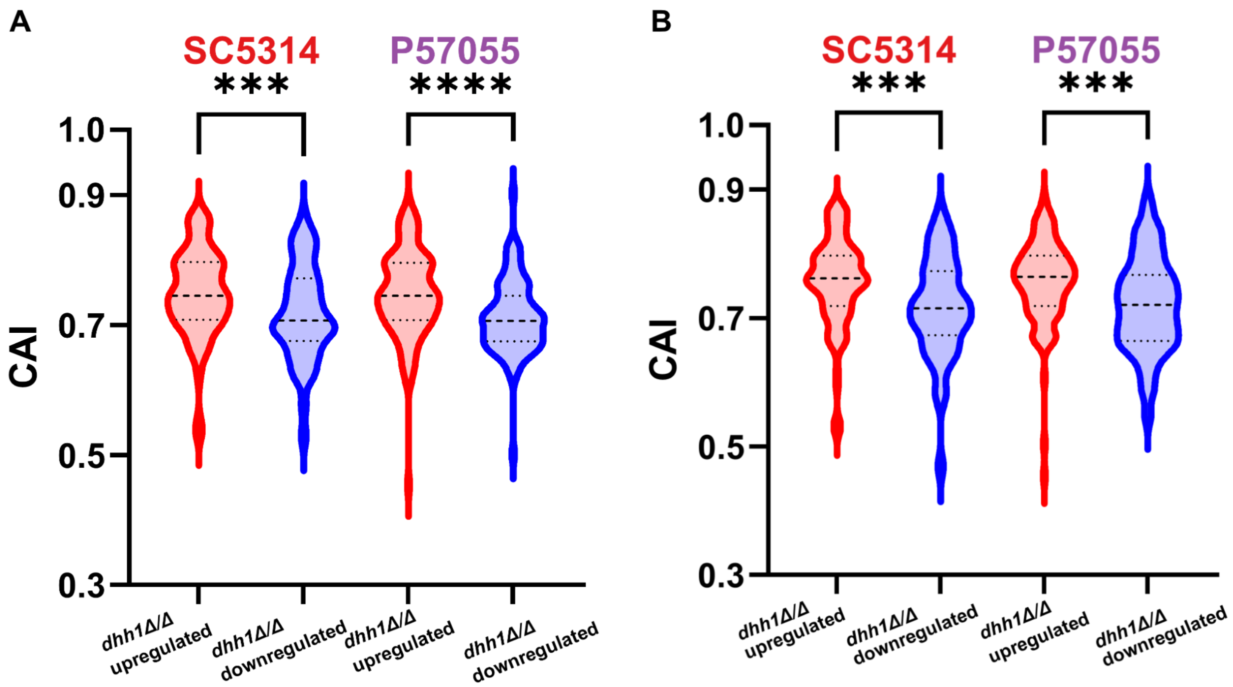


**Fig S6. Poor CAI is not associated with de-repression (upregulation) in *dhh1Δ/Δ*.** Upregulated genes in *C. albicans dhh1Δ/Δ* have higher CAI values than downregulated genes. CAIs of 100 most upregulated and downregulated genes in SC5314 *dhh1Δ/Δ* and P57055 *dhh1Δ/Δ* strains grown in **A.** YPD at 30°C and **B.** RPMI + 10% FBS at 37°C . (Mann-Whitney tests. ***P < 0., ****P < 0.0001).

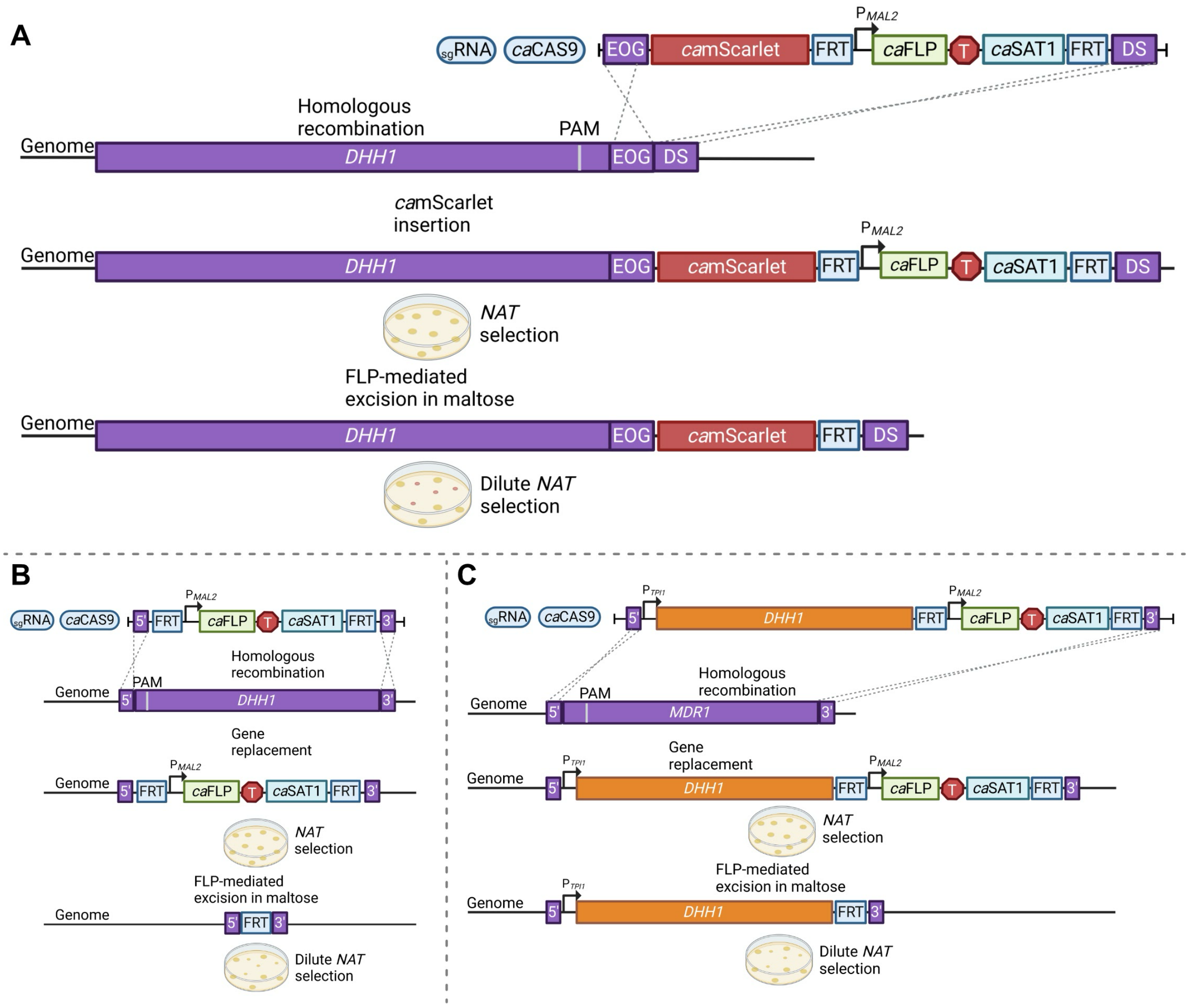


**Fig S7. Transient CRISPR-cas9 cloning diagrams.** Methods for **A.** inserting a fluorescent tag in the genome, **B.** deleting the entire ORF for a gene leaving only a 34bp FRT scar, and **C.** generating a *DHH1* complement at the *MDR1* site.

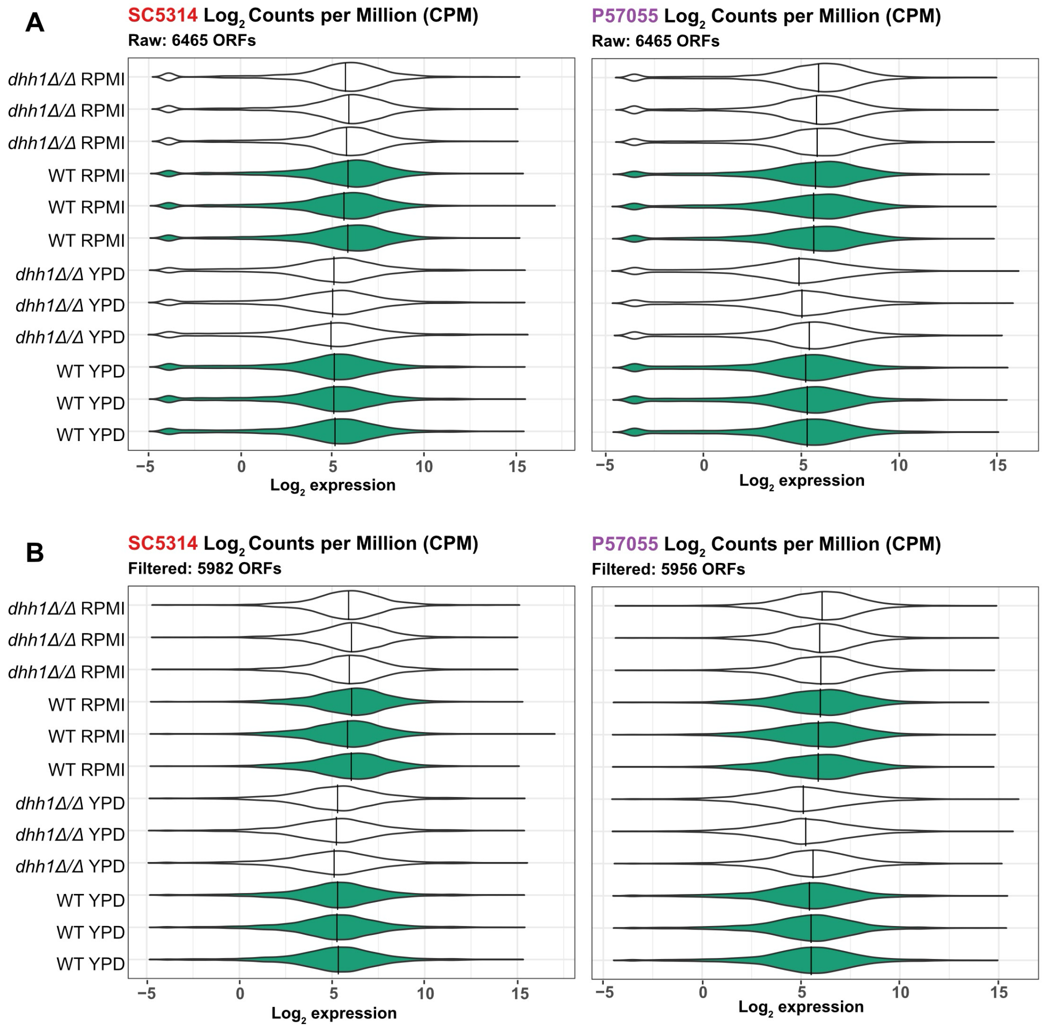


**Fig S8. Filtering out poorly transcribed ORFs in normalized RNAseq hits.** ORFs with less than one CPM hit in less than 3 experimental groups were removed prior to running DEseq2. **A.** Violin plots show raw and **B.** filtered hits for each strain and condition with ≥1 CPM in ≥ 3 experimental groups.

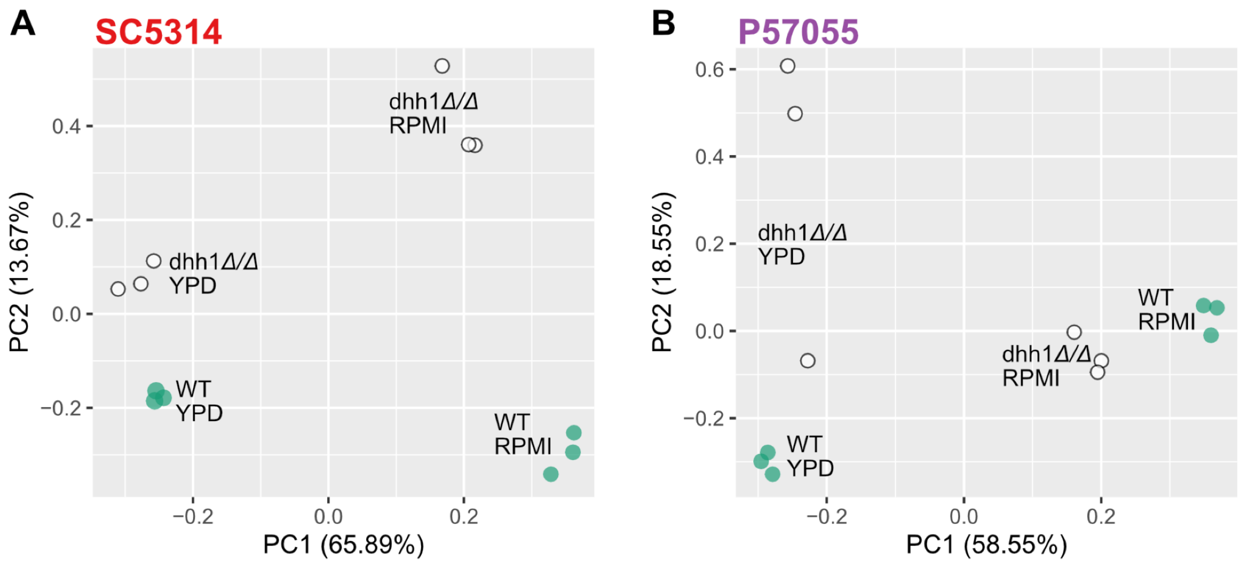


**Fig S9. Samples cluster by strain and growth conditions.** PCA plot displaying the filtered CPM from all 12 experimental groups clustering by condition in **A.** SC5314 and **B.** P57055.
